# Enhancing the Applied Force and Range of Axial Optical Tweezers

**DOI:** 10.1101/2023.04.21.537890

**Authors:** Zheng Zhang, Joshua N. Milstein

## Abstract

Axial optical tweezers provide a natural geometry for performing biomechanical assays such as rupture force measurements of protein binding. Axial traps, however, are typically weaker than their lateral counterparts requiring high laser power to maintain a well calibrated, linear restoring force. Here we show how to extend the spatial range over which well calibrated forces can be applied by considering aberration effects and extend the range of applied forces by accounting for the nonlinear response that appears when an optically trapped bead is moved far from the trap center. These refinements to the force calibration can be used to apply higher axial forces at reduced laser powers deeper into a sample. To illustrate the method, we reproduce both the linear extension regime and overstretching transition observed in dsDNA at significantly reduced laser powers.

http://dx.doi.org/10.1364/ao.XX.XXXXXX

---

Optical tweezers remain one of the most precise techniques for measuring forces involved in the interactions between biological molecules [1]. Axial optical tweezers, which apply forces perpendicular to surface tethered molecules (i.e., along the laser axis), may have certain advantages over more conventional trapping geometries [2]. Axial tweezers allow for the use of short tethers, which greatly reduces the compliance of the system and can significantly increase the signal-to-noise of a measurement. And compared to conventional dumbbell tweezers [1], axial tweezers are simpler to implement and the throughput is often much better as measurements can be quickly repeated on multiple tethered molecules. While an axial geometry seems straightforward enough, challenges in calibrating this technique, due to spherical aberrations and a focal shift that effect the location and strength of the trap as it is translated, have limited its adoption by the biophysical community. Yet, in recent years, these challenges have largely been overcome [2].

One crucial challenge that remains, however, is that at a given laser power, axial optical traps are typically much weaker than their lateral counterparts. Here, we show that by correcting for the nonlinear signal observed when a trapped microsphere moves far from the trap center, axial optical tweezers can apply calibrated forces significantly stronger than what could be achieved without this correction. By also considering signal aberrations deep within a sample, our method enables one to apply enhanced forces with weaker axial traps over a greater range of extension. This is illustrated by exploring both the linear force extension regime and the signature overstretching transition observed when pulling on dsDNA.

The optical setup, calibration of the sensitivity and trap strength with trap height, and a simple correction for the stage drift, have been presented in previous publications [2], so are provided here in the *Supplementary Materials*. To achieve a tight optical trap, we use an oil-immersion objective where the refractive index of the immersion oil matches the coverslip. However, the refractive index of the coverslip is higher than the aqueous trapping medium leading to refraction of the trap light (see *Supplementary Materials*). This induces a focal shift, which means that the height of the trap is not simply given by the relative stage displacement from the bead/surface touch point. To account for this, we make use of the observed interference between the laser light backscattered by the bead and then reflected forward by the coverslip. As shown [3], the system acts much like an optical cavity giving rise to a sinusoidal signal on our detector that can be fit by *S*(*X*)= *A* exp(*BX*) sin(*kL*_1_*X* + *ζ*)+ *P*(*X*), where *X* is the stage displacement, *A, B, L*1 and *ζ* are fit parameters, and *P*(*X*) is a second order polynomial. We have introduced the wave number *k* = 4*πn*/*λ*, where *n* is the refractive index of the trapping medium (*n ∼* 1.33) and *λ* is the wavelength of the trap laser (*λ* = 1064 nm). A fit is shown in Fig. 1A for the free bead signal at height *h* = *L*_1_*X* + *R* above the surface.

**Fig. 1.**
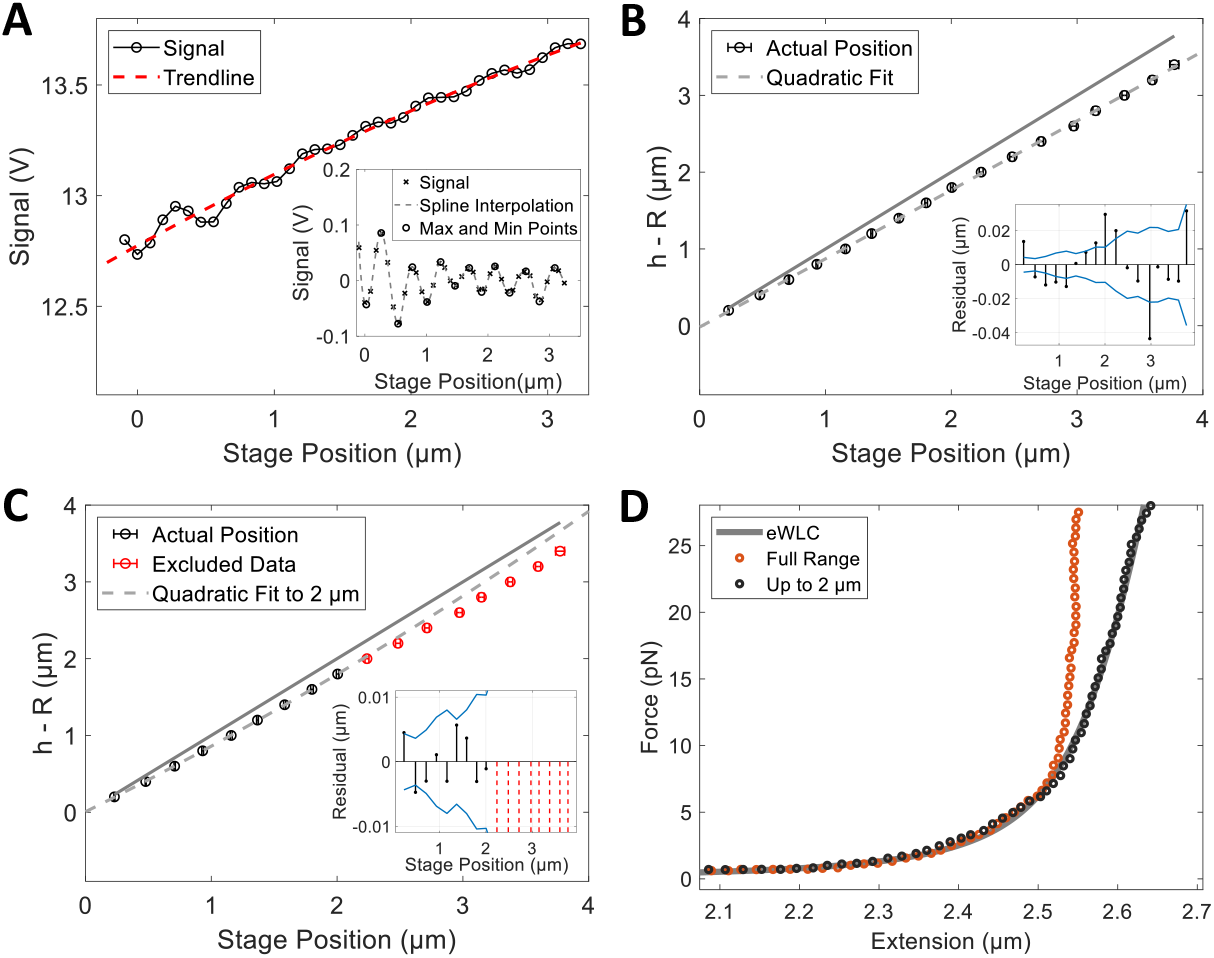
A) Signals recorded for an untethered bead relative to the coverslip. The red dashed line is a second order polynomial *P*(*X*) yielding a detrended signal (inset). B) Trap height (*h*) minus the bead radius (*R*) vs. stage position showing a quadratic fit to the whole range. C) Same as B for a fit up to stage positions of 2 *µ*m. For B and C, the solid line has a slope = 1, datapoints are an average of 27 signals, and error bars indicate standard errors. Insets show residuals to the fits with the two blue lines indicating the standard error of the data. D) Force-extension curves for 7782 bps dsDNA calculated from a fit over the full range (red circles) or restricted up to 2µm (black circles). The eWLC prediction is given by the solid line.

The above fit approximates the focal shift by a constant scaling factor *L*_1_, which can be further improved upon by introducing an additional quadratic term [4]. However, we find that when taking force-extension measurements of dsDNA at large extensions, it is extremely difficult to fit the signal to the degree of accuracy necessary to reproduce the known elastic properties of highly-stretched dsDNA, especially when the stage displacement *X* is large, which is necessary for large extensions. We next illustrate this point, showing how one can account for the focal shift to the necessary accuracy at large extensions.

We fit a second-order polynomial *P*(*X*) to the signal, use this fit to yield a detrended signal (i.e., *S*(*X*) -*P*(*X*)), then apply a spline interpolation to the data to find the maximum and minimum points (see inset of Fig. 1A).

These peaks, which should be separated by a distance of *λ*/4*n*, can be used as a ruler for defining the separation *h -R* between the bead and the coverslip surface. In Fig. 1B, we plot *h -R* at each peak as a function of the measured stage displacement. Note that the actual height of the bead is always less than the stage displacement. This curve simply quantifies the focal shift as we translate the stage.

Let us consider the results when we fit the curve with a second-order polynomial. Although the fit may appear to be adequate, we note that the residuals of the fit can be on the order of ∼ 20 nm or more (Fig. 1B). If we were to instead only fit the data up to stage displacements of 2 *µ*m, as opposed to across the full range of 4 *µ*m, the residuals to the fit within this range are now significantly lower (to within only a few nms), although the fit is quite poor at further distances (Fig. 1C). This suggests that the accuracy of a polynomial fit, at accounting for the focal shift, degrades as the trap is translated away from the surface. The accuracy with which we can measure the focal shift is critical to reproducing the force-extension of DNA at higher forces. When stretched, dsDNA initially behaves as a linear spring, but at increasing force the intrinsic elasticity of the polymer must be incorporated [5, 6]. Under typical buffer conditions, the worm-like chain (WLC) model holds (up to ∼ 10 pN) after which the response is essentially linear and may be described by the extensible worm-like chain (eWLC) model (up to ∼ 40 pN). Below this threshold, dsDNA remains in its B-form, but then the response curve begins to flatten and at ∼ 65 pN dsDNA begins to overstretch displaying a sudden increase in extension. The overstretched state depends upon subtle differences in ionic strength, temperature, and nucleotide sequence [7, 8] as well as on how force is applied to the DNA molecule. Under certain conditions this leads to a smooth transition while for others the force-extension can be highly stochastic.

In Fig. 1D, we show force-extension data for a 7782 kb dsDNA tether when we account for the focal shift with a second-order polynomial fit to all the signal data (as in Fig. 1B). While this calibration reproduces the WLC behavior of DNA, at further extensions, it significantly deviates from the predictions of the eWLC. This is because, within the steep linear region of the force-extension curve, small errors in the the position measurement translate into large deviations in the extension. We have found that including a third-order term in the polynomial expansion does little to improve the fit. Instead, we simply use an extrapolation of the fit restricted to data of stage displacements up to 2 *µ*m (Fig. 1C). Using this interpolation of the focal shift yields the force-extension curve shown in Fig. 1D, which now well agrees with the predicted eWLC behavior of dsDNA. We reproduced these results at two other tether lengths and the results are consistent (see *Supplementary Materials*). We suspect that the need to restrict the fit arises from artifacts associated with spherical aberrations at displacements beyond 2 *µ*m. Having improved the calibration for the trap height far from the coverslip surface, we now correct for the nonlinearity in the signal as the bead is displaced from the trap center. To illustrate the problem, we explore the force-extension curves observed in dsDNA under high applied forces. Figure 2A displays a series of force-extension curves for a 4.6 kbp dsDNA tether taken at laser powers ranging from 1-3 W out of the laser source. In our setup, this translates to ∼ 460 mW, 680 mW, and 1000 mW, respectively, at the trap. At both 680 mW and 1000 mW, the overstretching regime is clearly visible, but at 680 mW it appears to occur at a much lower applied force (∼ 50 pN). At an even lower laser power of 460 mW, the force-extension curve begins to flatten, deviating from the eWLC model, at just ∼ 25 pN.

**Fig. 2.**
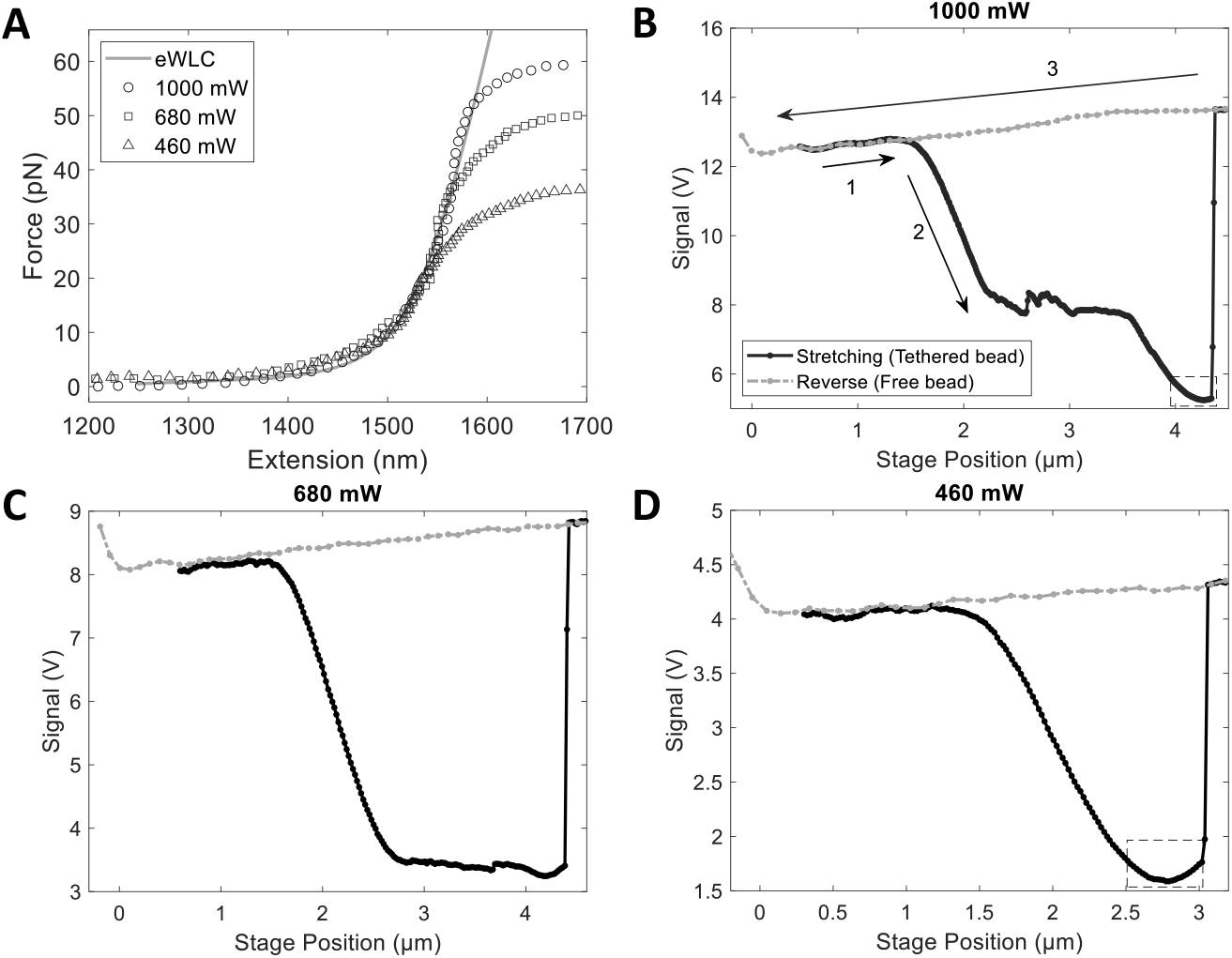
A) Force-extension at 1000 mW, 680 mW, and 460 mW of laser power. Measurements taken at less than 1000 mW appear to overstretch prematurely. A fit to the eWLC is shown (solid line). B) Photodiode signal vs. stage position during a measurement at 1000 mW. A tethered bead is first pulled away from the surface by moving the stage. Initially, the stage moves without affecting the trapped bead (1), but then the effects of the tether become apparent and the DNA responds like an eWLC, eventually, overstretching (2). Here, we reach a second linear regime after overstretching, just before rupture of the tether. After rupture, the stage is moved back to its initial position and the free bead signal is recorded (3). Signal vs. stage position at C) 680 mW and D) 460 mW. In both B and D, the nonlinear response of the trap is indicated by the dashed boxes, but is difficult to detect in C.

Figures 2B-D clarify the origins of this behavior. Here we display the photodiode signal as a function of stage position. The solid lines represent a dsDNA tethered bead displaced vertically by the trap until the tether ruptures, and the dashed-dotted lines are the signal from that same bead, now untethered, as the stage is scanned back across the range. The sawtooth pattern in the overstretching region is due to stepwise unpeeling of the dsDNA (Figs. 2B,C). To translate these measurements into the force-extension curves shown in Fig. 2A, we assumed that the applied force was linearly proportional to the displacement *x* of the bead from the trap center and that *x* was linearly proportional to the diode voltage. The proportionality constants are given by the spring constant *κ* and sensitivity *β*, respectively, which were measured at each axial position during the reverse scan.

When performing force-extension measurements at low laser power (460 mW, Fig. 2D), we observe a nonlinearity in the diode signal within a region where we expect the signal to be linear. As the stage is moved away from the trap and the DNA is stretched, the signal becomes linear in agreement with the eWLC, but then is observed to turn over. This same nonlinear behavior is seen at high laser power (1000 mW, Fig. 2B), but is not observed until the linear regime beyond the overstretching transition. However, when this nonlinear response in the signal occurs close to or within the overstretching regime, it is less obvious since the actual force required to extend the DNA tether is also beginning to flatten becoming stochastic about a relatively constant value, which obscures the signature shape of the nonlinearity (Fig. 2C). Ostensibly, these observations indicate that, for our setup, we need approximately 1000 mW of laser power to reach the overstretching regime. However, the force-extension curves at both 1000 mW and 680 mW appear qualitatively similar. The overstretching regime simply appears at too low an applied force when measured at 680 mW. This implies that there is something wrong in our force calibration at these lower powers that, if corrected for, would enable us to employ a lower laser power to access the overstretching regime.

The error arises because, upon extension of the DNA, the bead is dragged far from the center of the trap so that the signal *S* is no longer linearly proportional to the trap offset *x*. We could correct for this if we knew the functional form of *S*(*x*) far from the trap center. To obtain *S*(*x*), we note that within the linear region of the force-extension, pulling on the tether by displacing the stage drags the bead a similar distance across the trap with little further extension of the DNA. So long as Δ*x ≈*Δ*X*, where Δ*X* is the relative stage position, we can effectively map out *S*(Δ*x*) ≈*S*(Δ*X*). We recalibrate the force extension measurements, by correcting for the nonlinearity, based on this observation. Note, throughout, we will assume that the trap stiffness *κ* is still valid despite these large displacements from the trap center. This assumption will be justified by the accuracy with which we are able to reproduce the known polymeric properties of stretched dsDNA upon rescaling only the signal.

Let us take a closer look at the characteristic shape of the signal nonlinearity. In Fig. 3A, we obtain a signal (S) vs. stage displacement (X) curve obtained when stretching dsDNA at low laser power (60 mW). A nonlinear response is clearly seen at increasing stage displacement where the signal should remain linear. We then fit a hyperbola to the signal fitting data from the start of the linear regime to the minimum of the the measured signal at (*X*_*m*_, *S*_*m*_). The form of the hyperbola fit is as follows:

**Fig. 3.**
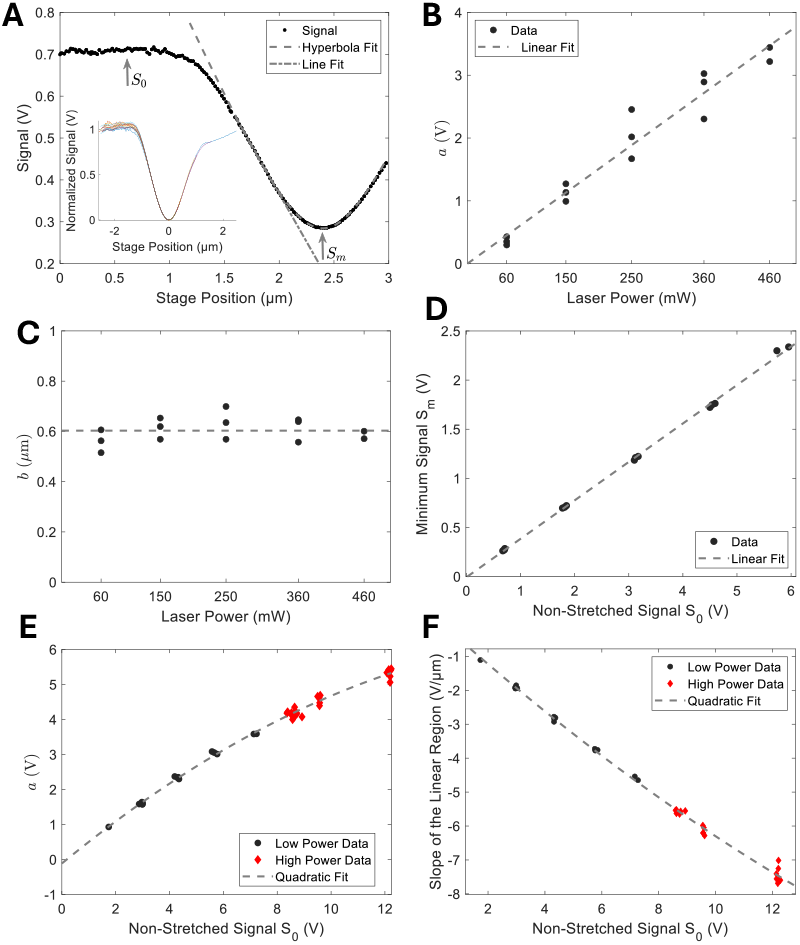
A) Low power signal displaying a hyberbolic inflection point at large stage displacements. *S*_0_ indicates the non-stretched signal and *S*_*m*_ the minimum of the nonlinearity. Inset: 14 signals from 60-460 mW centered at 0 and normalized by their *a* values. B) Measured values of *a* and C) *b* from the hyperbolic fits at low laser powers. The dashed line in (B) is a linear fit to the data and in (C) indicates the mean. D) *S*_*m*_ as a function of *S*_0_ is well fit with a linear function. E) The value of *a* and F) the slope of the linear region as a function of *S*_0_ at low power (black dots) and high power (red diamonds; 570-1000 mW). Dashed lines in (E) and (F) are quadratic fits.

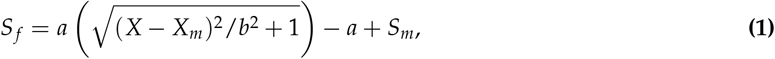

where *a, b, X*_*m*_ and *S*_*m*_ are fit parameters. This is repeated at increasing laser power (from 60 mW to 460 mW) so long as the hyperbolic shape of the nonlinearity is clearly observed within the linear force-extension regime. We find that the shape of the nonlinearity is maintained, it is just translated and scaled (inset to Fig. 3A).

The results of the fits for *a* and *b* are shown in Figs. 3B,C. We see that the parameter *b*, which describes the curvature of the nonlinear signal, remains relatively constant as we vary the laser power and that *a*, which scales the signal with laser power, appears to increase linearly. Similarly, the calibration curve in Fig. 3D, of the signal at the minimum of the hyperbola *S*_*m*_ as a function of the initial, “non-stretched” signal *S*_0_ (the points *S*_*m*_ and *S*_0_ are indicated in Fig. 3A), shows that for increased laser powers *S*_*m*_ is just linearly translated relative to *S*_0_. We next extend this fit analysis to higher laser powers to generate a final series of calibration curves for our force-extension measurements. Instead of formulating these curves as a function of laser power, which may drift during a series of measurements, it will be more convenient to plot them as a function of the readily accessible background signal *S*_0_. Fig. 3D shows *S*_*m*_ as a function of *S*_0_, which displays a strong linear trend. We will assume that this linearity holds, while also assuming that *b* remains a constant, and use these assumptions to simplify our fit to the data at higher laser powers. Again, we only fit the data within the linear region of the signal where *S*(Δ*x*) ≈*S*(Δ*X*) to yield Fig. 3E. The functional form of *a* is well fit by a quadratic function, which is not obvious at low powers (Fig. 3B). Finally, we plot the slope of the linear regime at each value of *S*_0_ (Fig. 3F). Figures 3D-F, and the constant value of *b* extracted from Fig. 3C, will enable us to correct the signal for an arbitrary force-extension measurement.

Now let us recalibrate data we obtained when overstretching dsDNA with 790 mW of laser power. When we displace the stage at a constant rate, we observe a linear response in the signal that flattens upon further extension, implying we have reached the overstretching regime (Fig. 4A). However, the force-displacement curve reaches the overstretching regime at only ∼ 55 pN (Fig. 4B) in comparison to our observation at 1000 mW where overstretching occurs closer to ∼ 65 pN (Fig. 4C) as expected. To correct the force-displacement curves, we first use Figs. 3D-F to extract the fit parameters *a* and *S*_*m*_ at the background signal value *S*_0_ (not shown) and the mean of Fig. 3C to obtain *b*. Note, the only remaining parameter in Eq. 2 is *X*_*m*_, but this is only an offset of the stage position and will not affect the shape of the hyperbola. We then generate a hyperbolic curve, representing the nonlinear response, shown overlaid on the data in Fig. 4A. We also either fit the linear portion of the data, or use the slope indicated in Fig. 3F, to extrapolate the linear region.

**Fig. 4.**
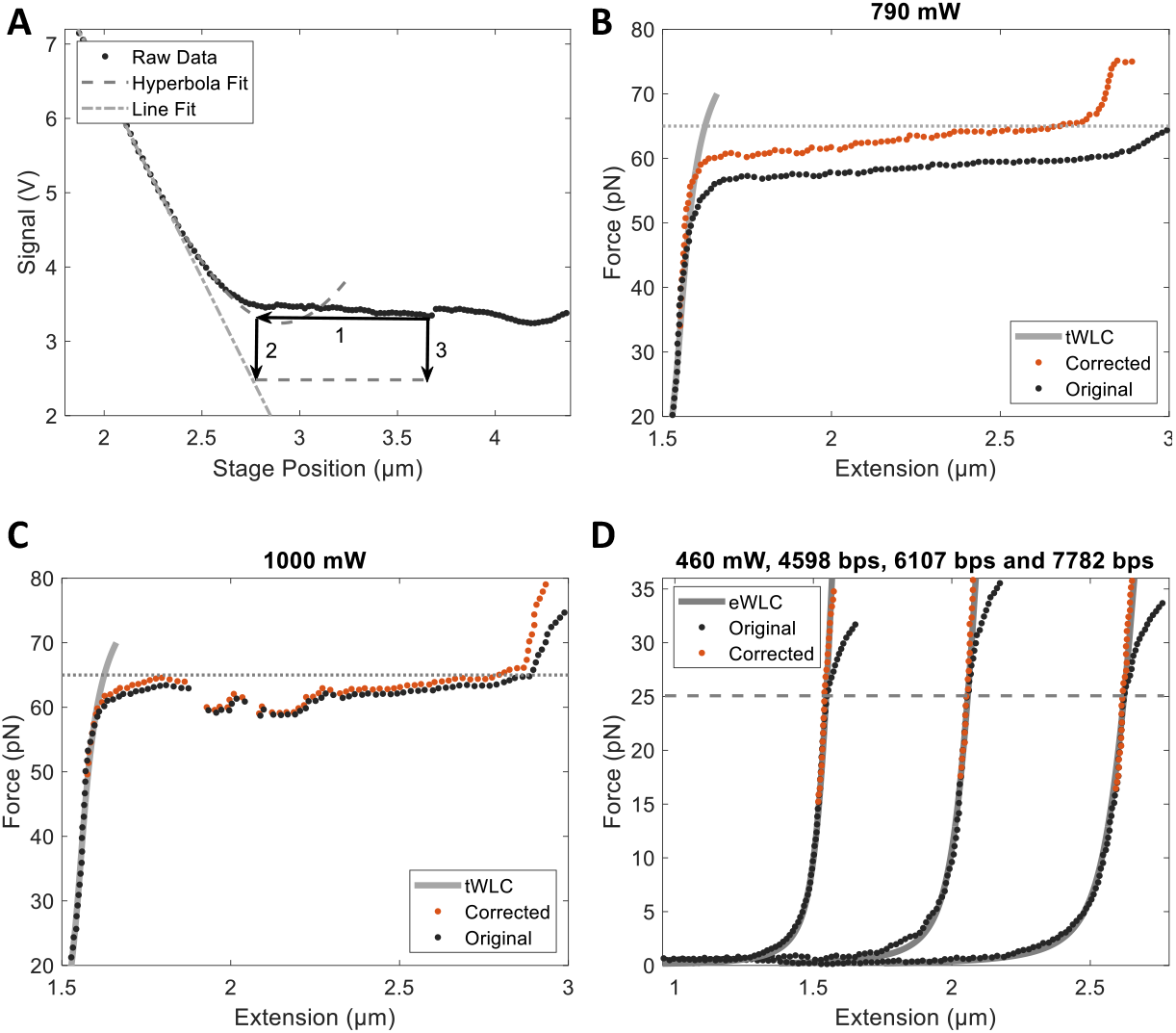
A) Signal vs. stage position. A hyperbola is fit to the data (dashed line) and the linear portion is extrapolated (dash-dotted line). Data points are first mapped onto this hyperbola then projected onto the linear fit. Force-extension curves at B) 790 mW and C) 1000 mW showing the raw data (solid lines) and corrected data (dash-dotted lines). The horizontal dotted line indicates 65 pN. Predictions of the tWLC are also shown (grey solid line). D) Correction result for three different lengths of dsDNA (left to right: 4598, 6107 and 7782 bps) compared with eWLC model at 460 mW. The dashed line indicates the position where the curves deviate from linearity.

The applied force can now be corrected by linearizing the signal. This is done by first mapping the measured signal onto points along the hyperbola and next mapping those points onto the extrapolation of the linear portion of the signal (Fig. 4A). Fig. 4B shows the results of the signal correction on the force-extension curve obtained at 790 mW. After rescaling, the curve now reaches the overstretching regime at a similar force as observed at 1000 mW (Fig. 4C). We also performed this rescaling on our data at 1000 mW of laser power (Fig. 4C). The resulting curve shows that even at 1000 mW, which is the maximum power our laser can attain, the signal still needs to be slightly corrected (although the deviation is only on the order of ∼ 1 pN). This signature behavior of overstretched DNA acts as a sanity check on our assumptions that went into the rescaling. We also provide a similar analyses for longer lengths of dsDNA and at lower laser powers (Fig. 4D). While this method enables a user to extend the range of forces accessible with axial optical tweezers, as the inflection point of the nonlinearity is approached, the error introduced by rescaling the signal increases significantly, limiting the range of correction. An error analysis on this rescaling is provided in the *Supplementary Material*. Here we extend the correction until the error is on the order of ∼ ± 3%, which translates to roughly ∼ ± 1pN at a maximum applied force of 35 pN. Even higher forces could be reached, at the current laser power, if we relaxed this criteria further, but the range of forces that can be applied by rescaling the signal is already increased by ∼ 1.5*x* (i.e., 25 pN to 35 pN) while introducing an error of only ∼ ± 1 pN at the highest accessible forces.

By accounting for aberrations and nonlinear effects discussed within this manuscript, axial optical tweezers can be used to explore a much greater range of forces, over longer distances, and at reduced laser powers. We hope this work will hasten the adoption of axial optical tweezers by the biophysical community, which may serve as a complementary and powerful single-molecule tool for exploring mechanobiology.

## Acknowledgments

We wish to thank Emiel Visser, Khadija Lakdawala and Sana Suboh for assistance at various stages of this work.

## Data Availability

Data underlying the results presented in this paper are not publicly available at this time but may be obtained from the authors upon reasonable request.

