## Supplementary Materials for "Enhancing the Applied Force and Range of Axial Optical Tweezers"

### 1. Axial force spectroscopy

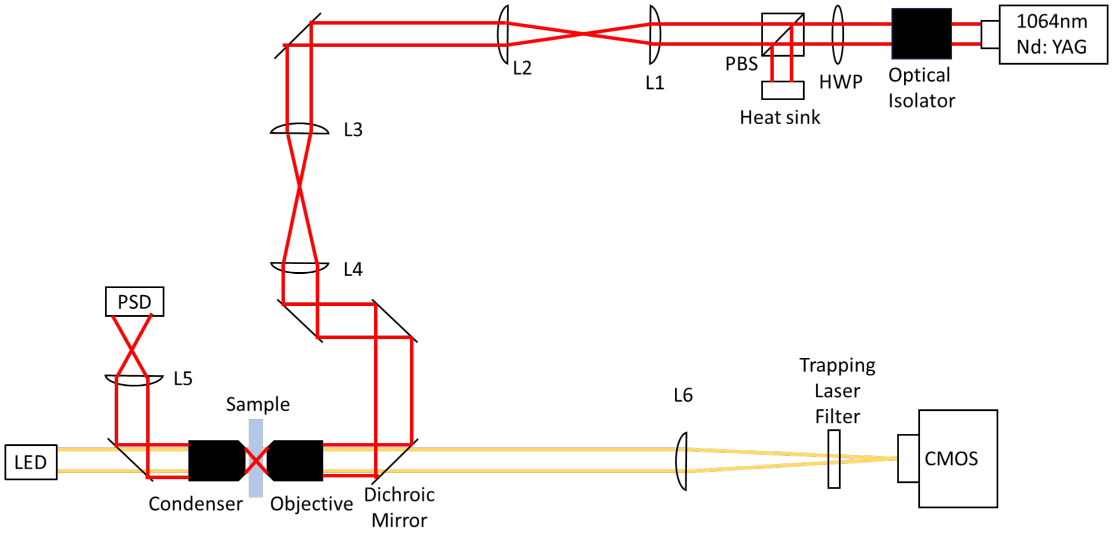


**Figure S1.1.** Optical setup of the axial optical tweezers.

The axial tweezers setup employs a standard configuration for single-beam optical tweezers. A Nd:YAG laser (Spectra Physics, 1064nm, BL-106C), used for optical trapping, is first passed through an optical isolator then circularized by passage through a pin-hole (not shown). A half waveplate (HWP), a polarizing beam splitter (PBS) and a heatsink are used together to change the power into the sample. Two sets of telescopes adjust the beam size (Olympus PLANAPO, 60x, oil-immersion, N.A.=1.42) to ~ 11 mm (full width at $1/e^2$ intensity), matching the size of the back aperture of the objective, to form a strong axial optical trap. A second objective (Olympus UPLFLN, 40x, N.A.=0.75) is used as a condenser to collect the laser light with its back focal plane conjugated by an additional lens onto a position sensitive diode (PSD, First Sensor, DL100-7-PCBA3). We used a red LED to illuminate the sample for bright field imaging with a CMOS camera (Pixielink PL-B781F). The sample chamber can be finely translated both axially (along the laser axis) and laterally (perpendicular to the laser axis) by a piezo-stage (Mad City Labs, Nano-LP100) adjusting the position of the optical trap relative to the coverslip surface. Note, the trap itself is held fixed in space. All data was acquired via LabView and processed by custom MATLAB scripts.

To apply well calibrated axial forces, we follow a procedure adapted from Mack et al (Mack et al., *Review of Scientific Instruments*, 2012). In brief, axial tweezers require a calibration for both the sensitivity and the trap strength at varying heights above the coverslip surface. This may be obtained by trapping a free bead within a fixed trap and varying the height by a vertical displacement of the stage. At each axial position, both the sensitivity and trap strength may be extracted from an analysis of the resulting power spectrum. The displacement of the bead from the trap center can then be obtained by comparing the observed intensity to the intensity of a free bead within the optical trap at that height. Accurately measuring the height of the trap center or of a trapped bead above the coverslip surface, while accounting for the focal shift (Fig. S1.2), is critical when performing force-extension measurements with axial optical tweezers. This, fortunately, can be precisely inferred from the backscattering of light between the trapped bead and the coverslip surface, which gives rise to an oscillatory signal as the trap is translated axially.

Instead of performing these calibrations in advance, in the measurements presented here on tethered DNA, we extend each construct by displacing the stage while recording the intensity until the tether ruptures. We then scan the now 'free' bead back toward the surface recording the sensitivity and trap strength at each height and using the resulting intensity scan to calibrate for the trap position. This approach is more robust than a single calibration step since sample drift can mostly be accounted for by simply aligning the backward scan with the initial, low-force portion of the forward scan. It is also a practical way to adapt the present technique to rupture force measurements that quantify protein or nucleic acid binding affinity.

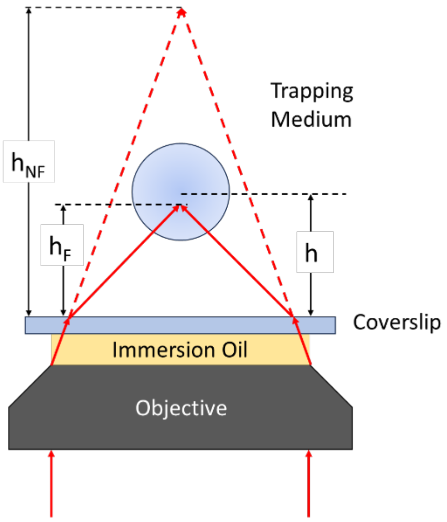


**Figure S1.2.** Illustration of the focal shift. A collimated laser beam is focused by the objective. The immersion oil and coverslip have a similar refractive index, which is higher than that of the aqueous trapping medium the bead is in. Due to this refractive index mismatch, the height of the laser focus is shifted from $h_{NF}$ to $h_F$ (not to scale). The actual trap height $h$ is then slightly downstream from $h_F$ because of the laser pressure.

### 2. Calibration of sensitivity and trap strength for force-extension measurements

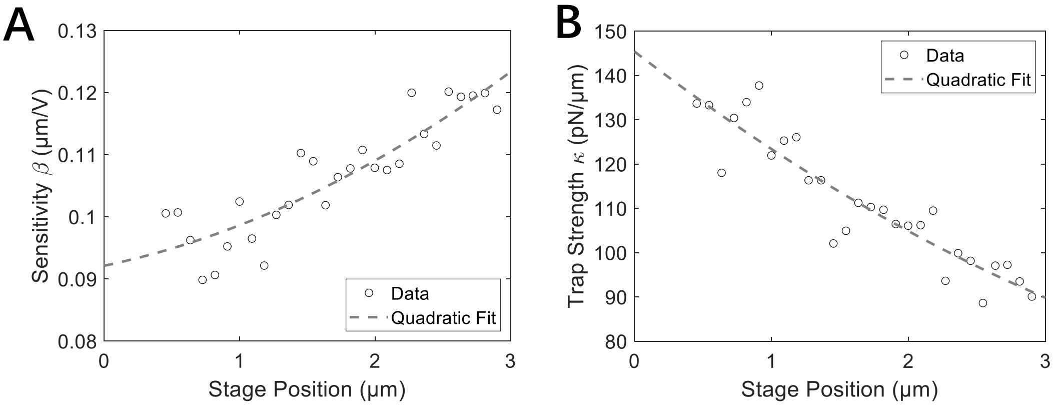


**Figure S2.** (A) The sensitivity $\beta$ and (B) trap strength $\kappa$ of the at different stage positions. Dashed lines result from a quadratic fit to the data.

With the optical trap remaining fixed in space, we move the stage until we trap a surface tethered bead. To center the bead in the trap, we first employ a low laser power (~60 mW, ~ 1 pN/μm trap strength) to pull the bead ~500 nm axially from the trap center. Then, we move the stage in the lateral direction to find the maximum intensity position. Lastly, we move the stage ~1 μm upward to relax the DNA.

After centering, we increase the laser current to the working laser power for our experiments. We keep the trap fixed and move the stage down, pulling the bead away from the trap center at a uniform speed. A voltage signal is recorded by a PSD as the stage descends, which will later be used to calculate the force-extension of the tethered DNA. We then continue until a rupture after which we reverse the stage direction moving it upwards (at ~ 100 nm/s) to retrieve the free bead signal. From the free bead signal, we can extract the sensitivity $\beta$ and trap stiffness $\kappa$ at each height from a fit to the power spectrum of the following form:

$$|P(f)|^2 = \frac{D_V}{\pi^2(f^2+f_c^2)}, \tag{S1}$$

where $D_V$ and the corner frequency $f_c$ are fitting parameters. The trap stiffness $\kappa$ and sensitivity $\beta$ are given by:

$$\kappa = 2\pi f_c \gamma, \tag{S2}$$

$$\beta = \sqrt{\frac{k_B T}{\gamma D_V}}, \tag{S3}$$

where $k_B$ is the Boltzmann constant, $T$ is the temperature, and $\gamma$ is the drag coefficient. Due to edge effects, the drag coefficient $\gamma$ is a function of bead radius $R$ at height $h$ above the coverslip surface. It can be approximated by Brenner's formula:

$$\gamma_\perp = \frac{\gamma_0}{1-\frac{9R}{8h}+\frac{R^3}{2h^3}-\frac{57R^4}{100h^4}+\frac{R^5}{5h^5}+\frac{7R^{11}}{200h^{11}}-\frac{R^{12}}{25h^{12}}}, \tag{S4}$$

where $\gamma_0 = 6\pi\eta R$ is the drag coefficient for a sphere of radius $R$ in an infinite medium of viscosity $\eta$. Figure S2 shows the axial position dependent behavior of the sensitivity $\beta(X)$ and trap strength $\kappa(X)$. Notice, the observed oscillations are an artifact arising from light backscattered between the bead and coverslip surface. In practice, as shown in the figure, we fit a quadratic function to the data to obtain $\beta(X)$, $\kappa\ (X)$ at all axial positions. The difference between the tethered bead signal and free bead signal $\Delta s\,(X)$ can then be used to calculate the force applied by the optical tweezers at each axial position as:

$$F(X) = \kappa(X) \cdot \beta(X) \cdot \Delta s(X), \tag{S5}$$

where $\Delta s\,(X)$ is the change in the measured signal.

### 3. Correcting for stage drift

When using axial optical tweezers, one of the biggest issues is the axial drift of the stage. Accurate measurement of the distance and the force largely relies on an accurate measurement of stage position relative to the contact point between the bead and the surface. Axial drift will lead to an inaccurate measurement of the stage position as we only record the stage displacement from the piezo stage controller. Figure S3.1 shows a pair of signals with and without drift correction. We can clearly see that the sinusoidal interference pattern doesn't initially match (Fig. S3.1B). Here we used DNA with a length of ~1.5 μm. After centering the tether, we move the stage up to a position where it is ~0.5 μm from the touch point before starting to record the stretching signal, and the signal here is clearly not at the expected position from the uncorrected signal.

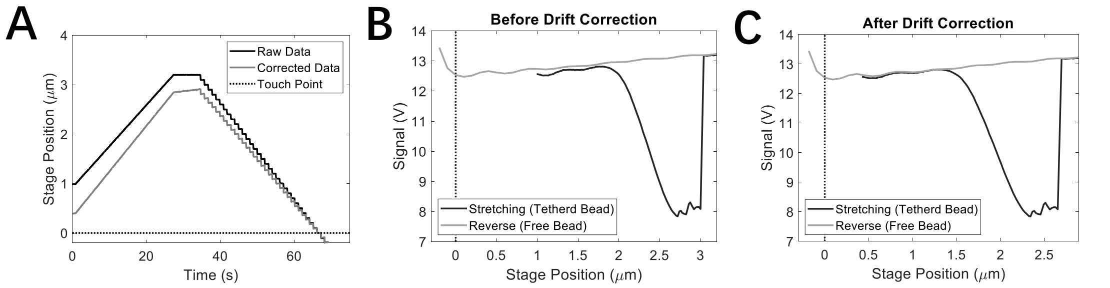


**Figure S3.1.** Correction of the axial stage drift. (A) Raw stage position and corrected stage position. The touch point is set to 0, and stage position is the position of the stage relative to the touch point. (B) Signals before and (C) after axial drift correction. The measurement starts at the left of the stretching signal. As the stage moves down, the signal decreases, the force increases, and the tether eventually breaks, which is the right part of the stretching signal. Then we scan the stage back until it touches the bead again (Reverse scan, right to left.).

There are many methods to correct axial drift of which a common one is using a reference bead stuck on the surface. However, this method requires complex image processing for each measurement. There are also commercial solutions to correct the drift. Fortunately, for our axial optical tweezers' setup, when the laser power is kept constant during one measurement, we found that the axial drift of the stage can be approximated by a linear drift with a constant speed (Figure S3.2) and can be corrected by incorporating a linear drifting speed $v_d$ through the equation:

$$S_c = S_0 + v_d t, \quad \text{(S6)}$$

where $S_c$ is the corrected stage position, $S_0$ is the raw stage position recorded from the piezo stage controller, and $t$ is time. One important thing to note is the rightmost points in both the stretching and reverse signal in Figure S3.1B can be regarded as identical as long as the time interval between the switch between the stretching and reverse signal is negligible. Signals after correction are shown in Figure S3.1C. After this simple correction, the sinusoidal interference patterns match each other well, and the starting point of the stretching signal is ~0.5 μm from the touch point, proving that our drift correction method works well. However, in practice, the precision of this method depends on the actual drift pattern because we employ a linear approximation. Also, when the sinusoidal pattern is not clear, it will be hard to match the two signals, so can care should be taken when employing this simple correction method.

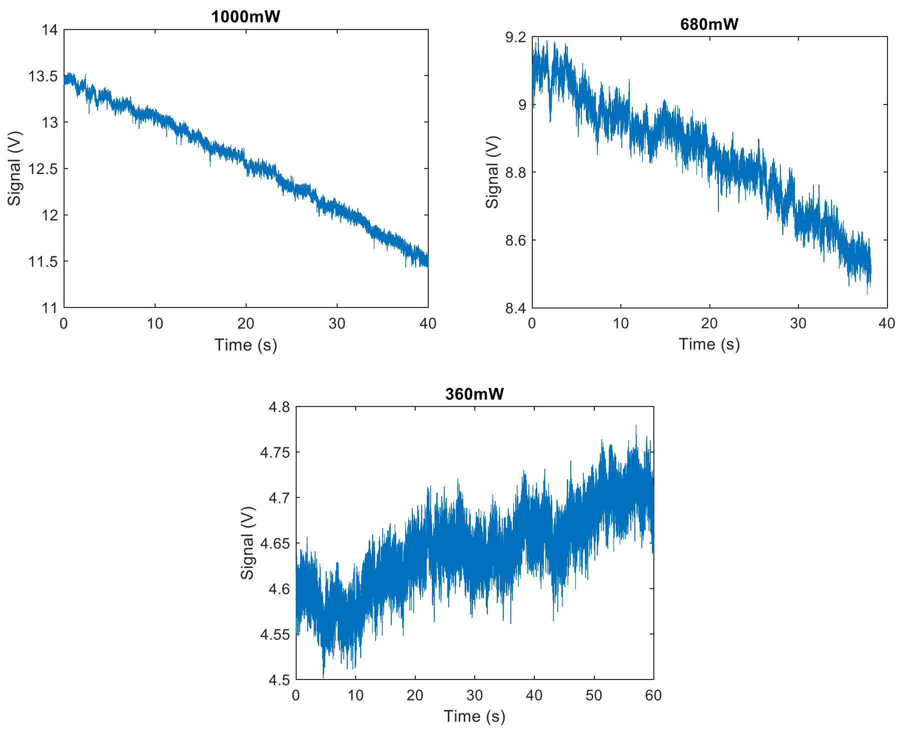


**Figure S3.2.** Signals from a bead stuck on the coverslip surface at 1000 mW, 680 mW, and 360 mW laser powers. The signal is approximately linearly proportional to the axial position of the optical trap.

### 4. Bead-coverslip separation calculated by two fitting methods

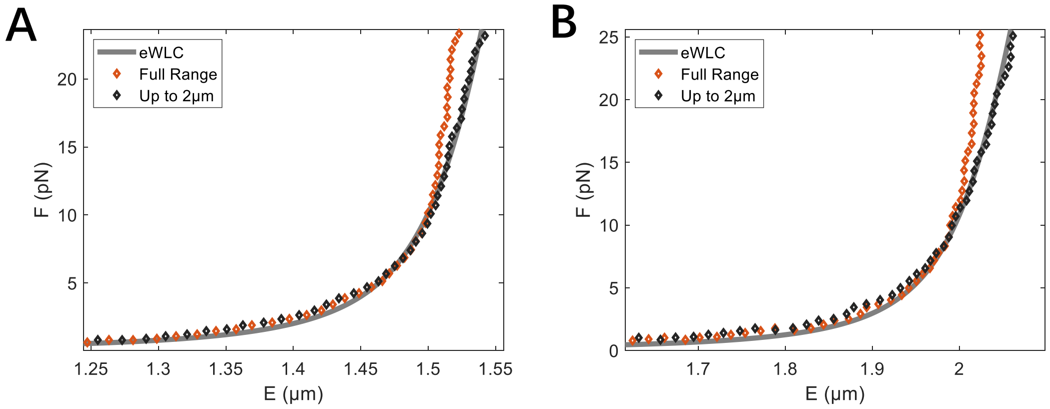


**Figure S4.** FE curves calculated using two methods. The length of the dsDNA is (A)6107 bps or (B)7782 bps.

### 5. Quantification of the error in the signal correction

Here we investigate the limits of correcting the nonlinearity in the signal. The predicted value of $S_m$ , which is the value of the signal nonlinearity at the inflection point, is the main source of error for the correction. This error is amplified as the measured signal is corrected increasingly closer to the inflection point. Figure S6A shows the error between the predicted $S_m$ from Figure 2D and the measured $S_m$ directly from the signal (for 24 signals) at low laser powers that cannot overstretch the dsDNA. The errors are, roughly, between ±4%. We investigated how this error in $S_m$ will affect the correction and the result is shown in Figures S6 B-D. The stage positions indicated in Figures S6 B and C are the position from the minimum signal ($X_m$, $S_m$). The position $X_m$ is set at zero. As we can see, the error in the force prediction gets larger as the stage position approaches the minimum signal.

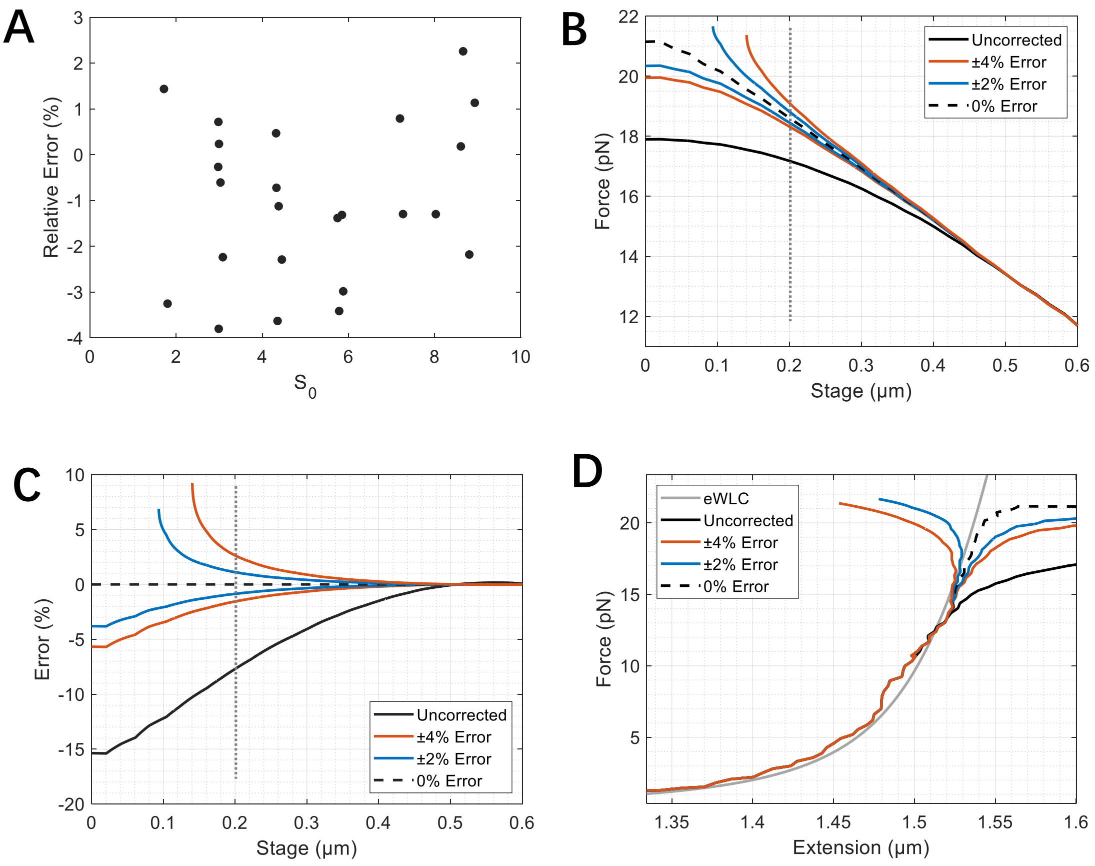


**Figure S6.** (A) Relative errors between predicted $S_m$ from Fig. 2D and $S_m$ obtained by directly fitting the signal (for 24 signals at different laser powers). Different background signal values $S_0$ correspond to different laser powers (see Fig. 2 caption). (B) Predicted forces at different stage positions from uncorrected signals and corrected signals with varying error in $S_m$. The dotted line indicates a cutoff of 0.2 μm from the inflection point of the nonlinearity. (C) Relative error in the corrected force for various levels of error in $S_m$ at different stage positions. The dotted line indicates a 0.2 μm cutoff. (D) Force-extension curves calculated from uncorrected signals and corrected signals with different errors in $S_m$.

Here we set 0.2 μm as an arbitrary cutoff before the error gets too large. This stage position corresponds to an 8% force correction comparing the uncorrected signal and the 0% error signal. At this position, the displacement from the trap center is ~820 nm before the correction and is ~910 nm after the correction. This is far from the trap center, yet our method yields the correct overstretching forces for dsDNA at this trap displacement (Figs. 4 B,C). Note, one might be concerned that the restoring force of the optical trap is no longer linear either, but if this were the case than the force we measured should be much larger than the theoretical value because the trap strength should decrease, but we don't see this effect in our results.

We can see from Figures S6 B-C that the non-linear effect starts to be significant at ~0.5µm, and the force is 14 pN (laser power is 250mW for this signal). After our correction, at ~0.2µm, the force is ~19 pN, which means that we can access 36% more of the force range compared with when the non-linear effect is not corrected.
